# An ammonium transporter is a non-canonical olfactory receptor for ammonia

**DOI:** 10.1101/2021.03.31.437861

**Authors:** Alina Vulpe, Hyong S. Kim, Sydney Ballou, Shiuan-Tze Wu, Veit Grabe, Cesar Nava Gonzales, Silke Sachse, James M Jeanne, Chih-Ying Su, Karen Menuz

## Abstract

Two families of ligand-gated ion channels function as olfactory receptors in insects. Here, we show that these canonical olfactory receptors are not necessary for responses to ammonia, a key ecological odor that is attractive to many insects including disease vectors and agricultural pests. Instead, we show that a member of the ancient electrogenic ammonium transporter family, Amt, is a new type of olfactory receptor. We report two hitherto unidentified olfactory neuron populations that mediate neuronal and behavioral responses to ammonia. Their endogenous ammonia responses are Amt-dependent, and ectopic expression of either *Drosophila* or *Anopheles* Amt confers ammonia sensitivity. Amt is the first transporter known to function as an olfactory receptor in animals, and its role may be conserved across insect species.

## Introduction

Insect ORNs detect odors with ligand-gated ion channels, nearly all of which are members of the odorant receptor (OR) and ionotropic receptor (IR) families [1–5]. These receptors define the odor tuning of individual olfactory receptor neurons (ORNs), and the stereotyped receptor combinations expressed by neighboring ORNs define functional subtypes of sensilla, sensory hairs found on the antenna. In *Drosophila*, the best characterized insect model, the sensillar localization and odor response profiles of most OR and IR receptors has been characterized [5–7].

Numerous hematophagous insects including mosquitos, sand flies, lice and triatomine bugs are attracted to ammonia, a kairomone released in human sweat and breath [8–10]. Low levels of ammonia also attract non-biting insects such as the genetic model organism *Drosophila* and several species of tephritid flies, important agricultural pests [11, 12]. Its effectiveness as an attractant has spurred the use of ammonium-containing solutions in commercial insect traps [9–11]. In every insect species examined, ammonia activates ORNs housed in sensilla with a grooved peg morphology [9], also known as known as coeloconic sensilla. The molecular basis of ammonia sensing has been primarily examined in *Drosophila*, in which robust responses to low levels of ammonia (NH_3_) are observed in the ac1 subtype of coeloconic sensilla [7, 13, 14], and several studies have linked this response to the Ir92a receptor expressed in one of the three identified ac1 ORNs [7, 12, 15]. Given the widespread importance of ammonia to insect behavior, it is surprising that the genomes of most insects lack Ir92a orthologs [1].

Here we uncover previously unidentified ORNs and demonstrate that they mediate both neuronal and behavioral responses to ammonia. Surprisingly, these ORNs do not detect ammonia using a conventional IR or OR receptor, but rather utilize a highly conserved ammonium transporter as a novel type of olfactory receptor.

## Results

### Ammonium transporters in a new ORN class

We previously found that ammonia detection in *Drosophila* ac1 sensilla depends on Amt [14], a member of the conserved Amt/Rh/MEP ammonium transporter family with members in bacteria, fungi, plants, and animals [16, 17]. The ortholog in *Anopheles* mosquitoes, AgAmt, can restore ammonia responses in Amt mutant flies [14] and supports the typical ammonium-selective inward current found in members of this transporter family when heterologously expressed [18]. The precise reason why Amt is indispensable for ammonia detection remains unclear. In *Drosophila*, Amt expression is highly enriched in the antenna, where it was detected in support cells surrounding neurons in ac1 sensilla and the sacculus [14].

Apart from Amt, insect genomes contain a second ammonium transporter, Rh50, which has widespread expression in multiple tissues [16, 18]. Within the antenna, Rh50 transcript level is reduced ∼10-fold in *atonal* flies that selectively fail to develop coeloconic sensilla (Figure S1A) [14]. This observation suggests a potential connection between Rh50 and ammonia detection by coeloconic sensilla. We therefore generated a Rh50 reporter line to identify Rh50^+^ cells within the antenna. The scattered population of labeled cells had a neuronal morphology and was positive for the neuronal marker elav (Figures 1A and 1B). The reporter line was faithful because all antennal GFP^+^ cells in *Rh50-GAL4; UAS-GFP* flies were labeled by a Rh50 anti-sense probe (Figures S1B-S1D). Thus, Rh50 expression in the antenna was found solely in a subset of neurons.

**Figure 1.**
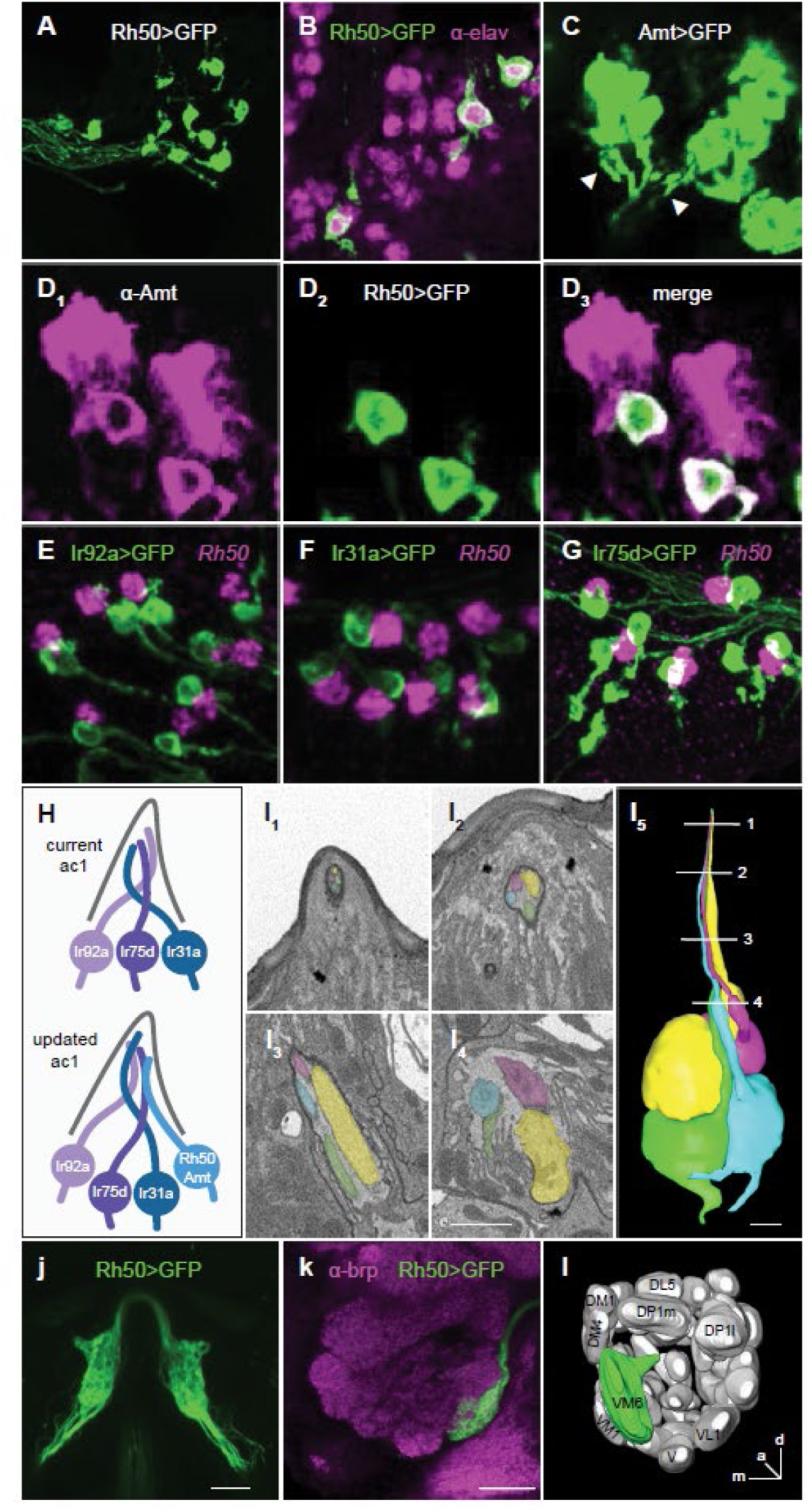
Ammonia transporters mark the expression of a previously unidentified ac1 ORN. (A) Antennal section from a *Rh50>GFP* fly stained with anti-GFP antibody. (B) *Rh50>GFP* fly antennal section stained with anti-GFP (green) and anti-elav (magenta), a marker for neurons. (C) High gain confocal image of an *Amt>GFP* fly antennal section stained with anti-GFP. Labeled axons (arrowheads) are emerge from clusters of GFP^+^ cells. (D) Immunostaining with anti-Amt (magenta, D_1_) and anti-GFP (green, D_2_) on antennal sections from *Rh50>GFP* flies. D_3_, Rh50^+^ neurons co-express Amt. (E-G) Antennal sections from *Ir92a>GFP* (E), *Ir31>GFP* (F), and *Ir75d>GFP* (G) flies labeled with an *in situ* hybridization probe for Rh50 (magenta) and an antibody against GFP (green). (H) One of three known ac1 ORNs is labeled in each line. One Rh50^+^ neuron consistently neighbors each GFP^+^ neuron (E to G), indicating that Rh50 is found in a 4th ac1 ORN (H). (I) Serial block face electron microscopy images of a coeloconic sensillum with four ORNs, each labeled a different color. A three dimensional reconstruction of the four ORNs is shown in (I_5_). Numbered lines indicate the locations of individual sections shown in (I_1-4_). Scale bars, 1 µm. (J) Two photon *in vivo* image of the bilateral antennal lobe glomeruli innervated by Rh50^+^ axons in a *Rh50>GFP* fly. Scale bar, 20 µm. (K) Confocal image of an antennal lobe from a *Rh50>GFP* fly brain immunolabeled with antibodies targeting GFP (green) and brp (nc82, magenta), a neuropil marker used to delineate glomeruli. Scale bar, 20 µm. In both J and K, glial GFP expression driven by *Rh50-GAL4* was suppressed with *repo-GAL80* [39] to improve visualization of the ORN projections. (L) Diagram of the ventromedial location of the glomerulus innervated by Rh50^+^ ORNs, which corresponds to VM6.

Might Amt likewise be expressed in olfactory neurons in addition to its known expression in support cells? Indeed, close examination of antennal sections from *Amt-GAL4; UAS-GFP* flies revealed weak GFP^+^ axons emerging from strongly labeled support cells in ac1 sensilla (Figure 1C). A similar antennal expression pattern of an AgAmt reporter line was recently reported in mosquitos [19]. Immunohistochemistry with an anti-Amt antibody revealed that Amt is co-expressed in Rh50^+^ neurons (Figure 1D). Consistent with the reported antennal expression pattern of Amt, Rh50^+^ ORNs were found only in ac1 sensilla, but not in the other three classes of surface coeloconic sensilla (Figures S1E-S1H) [15].

The three reported neurons in ac1 sensilla express Ir92a, Ir31a, and Ir75d olfactory receptors, respectively [15]. To determine which ac1 ORN expresses the two ammonium transporters, we examined Rh50 expression with *in situ* hybridization on antennal sections from flies in which one of the three ac1 neurons was labeled by a transgenic reporter. Surprisingly, the Rh50^+^ neurons did not co-localize with any of the three known neurons, but instead always neighbored the labeled ac1 ORNs (Figures 1E-1G). This indicated that Rh50 and Amt are co-expressed in a previously undetected fourth neuron in ac1 sensilla (Figure 1H).

Early electron microscopy work only identified *Drosophila* coeloconic sensilla that house two or three neurons [20]. We therefore examined recent antennal serial block-face scanning EM (SBEM) datasets [21, 22] to resolve this discrepancy, and found that approximately one in four coeloconic sensilla do in fact contain four ORNs (Figure 1I; Figure S1I). Further, coeloconic sensilla with two, three, and four neurons are unevenly distributed over datasets acquired from distinct antennal regions (Figure S1J), which could explain why coeloconic sensilla with four neurons were missed in the prior study.

The axonal projections of all ORNs expressing the same receptor coalesce into a bilateral pair of glomeruli in the antennal lobe. Examination of *Rh50>GFP* flies revealed that the terminals of Rh50^+^ ORNs converge to a large ventromedial glomerulus (Figures 1J-1L). This glomerulus corresponds to the previously observed “orphan” glomerulus VM6 [5, 23], recently shown to be targeted by neurons sharing a similar developmental origin as the other ac1 ORNs [23–25]. Because VM6 is difficult to discern with neuropil staining, it has been the source of confusion in recent antennal lobe atlases, where it was merged with VP1 [26] or listed as VC5 [27, 28].

### Amt/Rh50^+^ ORNs respond selectively to NH_3_

The identification of the previously undetected Amt/Rh50^+^ ac1 ORN raised the question of its odor response profile. Previous work screening ac1 sensilla with >150 odorants identified a handful that elicit neuronal spiking responses [7, 13]. Although responses to ammonia and many amines were previously ascribed to the Ir92a^+^ ORN [7, 12, 15], we wondered whether the fourth ac1 ORN may mediate a portion of those responses. To address this question, we examined odor responses through calcium imaging of the antennal lobe in *Rh50>GCaMP7* flies. Ammonia reliably evoked large calcium responses, whereas there was no response to water or any of the tested amines (Figure 2A). In contrast, GCaMP imaging of Ir92a^+^ ORNs revealed that these neurons respond broadly to ammonia and to amines (Figure 2B), consistent with previous reports [7, 12]. Neither of these neurons is likely to detect alkaline pH, which may result from ammonia application, because ac1 neurons were unresponsive to two basic amines at 1% concentration, butylamine (pKa 10.8) and isoamylamine (pKa 10.6), even though 0.01% ammonia (pKa 9.4) induced robust responses (Figure S2). Thus, Amt/Rh50^+^ ORNs are selectively tuned to ammonia.

**Figure 2.**
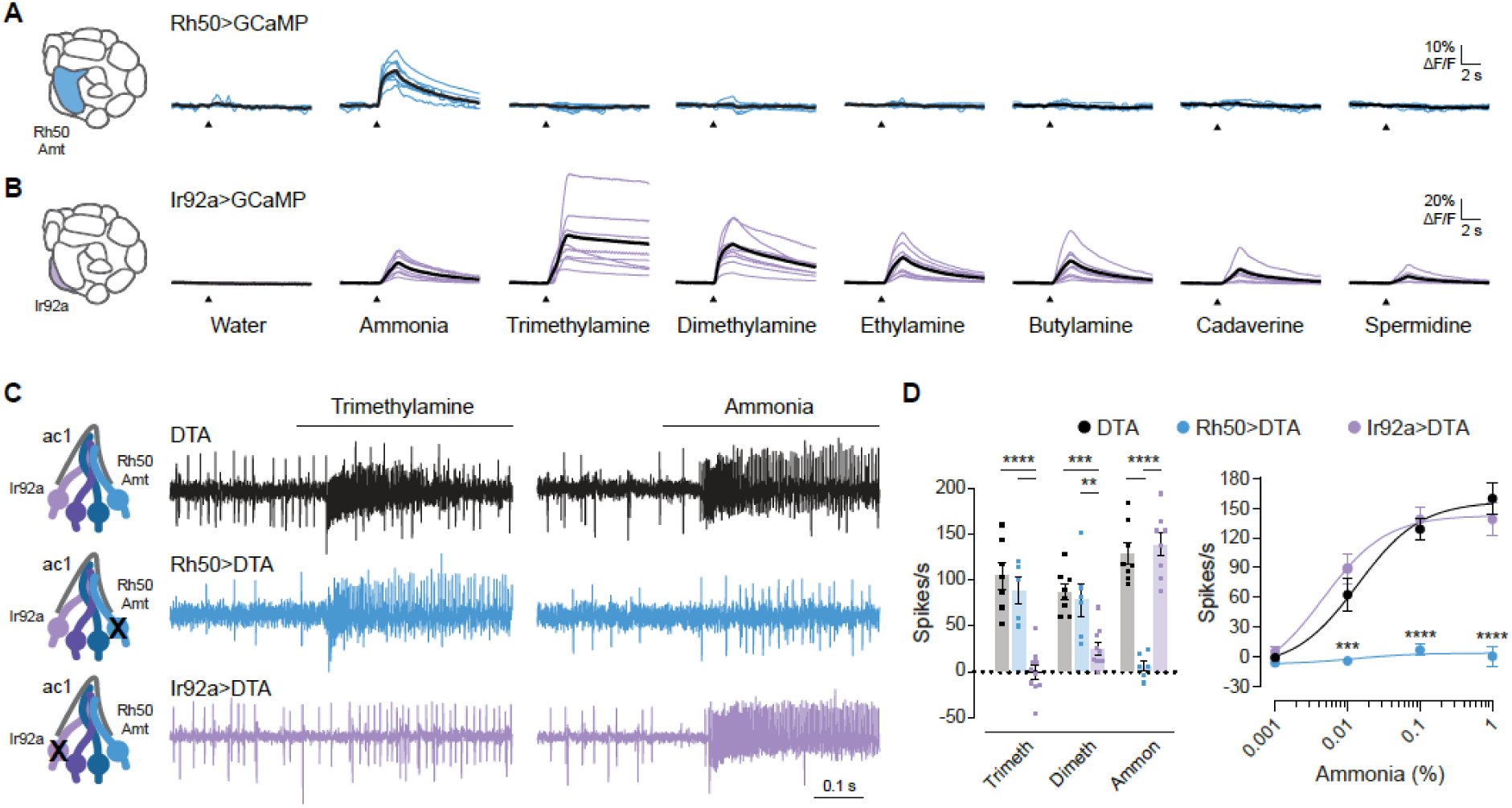
Amt/Rh50^+^ ORNs selectively respond to ammonia. (A) Antennal lobe calcium responses to water, ammonia, and several amines in axonal projections labeled in *Rh50>GCaMP7s* flies. All odors were delivered at concentration of 0.01%. Blue lines are responses in individual flies, and black lines are mean responses. (B) Antennal lobe calcium responses in axonal projections labeled in *Ir92a>GCaMP7s* flies. Purple lines are responses in individual flies, and black lines are the mean responses. (C) Representative traces of extracellular recordings of action potentials elicited by 1% trimethylamine and 0.1% ammonia in ac1 sensilla in which diphtheria toxin (DTA) was used to ablate Rh50^+^ ORNs (blue) or IR92^+^ ORNs (purple). *UAS-DTA* flies were used as a control (black). (D) Left, quantification of odor responses in *UAS-DTA* (black), *Rh50>DTA* (blue), and *Ir92a>DTA* (purple) flies (n= 5-10 sensilla). Right, dose-response curve of responses to increasing concentrations of ammonia in *UAS-DTA*, *Rh50>DTA*, and *Ir92a>DTA* flies (n= 6-8 sensilla). The dose-response data for 0.1% ammonia are replotted in the bar graph to show individual data points.

### Amt/Rh50^+^ ORNs mediate spiking responses to NH_3_

Given that both Amt/Rh50^+^ and Ir92a^+^ neuronal terminals showed ammonia-sensitivity with GCaMP imaging (Figures 2A and 2B), we next examined the relative contributions of these neurons to ammonia-induced spiking in ac1 sensilla. There is a nonlinear relationship between GCaMP7s responses and action potential firing because this highly sensitive calcium sensor signal can detect single action potentials but saturates at low firing frequencies [29]. Although several studies have ascribed ammonia induced spiking in ac1 sensilla to the Ir92a receptor and associated ORN, this was inferred indirectly from GCaMP imaging data demonstrating the ammonia-sensitivity of Ir92a^+^ ORNs and the ammonia-insensitivity of the two other previously known ac1 ORNs [7, 12].

We recorded from flies in which either the Ir92a^+^ or Amt/Rh50^+^ neurons were ablated through the selective expression of diphtheria toxin. Ablation of the Amt/Rh50^+^ ORNs abolished the large amplitude action potential responses in ac1 sensilla over a broad range of ammonia concentrations, whereas loss of Ir92a^+^ ORNs had no significant effect (Figures 2C and 2D). In contrast, ac1 spiking responses to trimethylamine and dimethylamine, odorants that strongly activate Ir92a^+^ ORNs in calcium imaging [7, 12] (Figure 2B), were unaffected by the loss of Amt/Rh50^+^ ORNs but were nearly eliminated by Ir92a^+^ ORN ablation (Figures 2C and 2D). Consistent with these data, an Ir92a loss-of-function mutant had similar effects on ac1 odor responses as Ir92a^+^ ORN ablation (Figure S3). Thus, Ir92a^+^ ORNs function primarily as amine detectors, whereas the Amt/Rh50^+^ ac1 ORNs mediate the robust spiking observed in response to ammonia.

### Sacculus Amt/Rh50^+^ ORNs also respond to NH_3_

Examination of *Rh50>GCaMP* antennae revealed a second population of Rh50^+^ neurons in the sacculus (Figure 3A, arrowhead), a three-chambered cavity invaginated from the antennal surface [30]. This was reminiscent of Amt expression, which is expressed in sacculus chamber III in addition to ac1 sensilla [14]. There were 26+/-3.6 (SD, n=10) Rh50^+^ ORNs when examined in whole-mount *Rh50>GFP* antennae, suggesting that each of the previously reported 22-26 chamber III sensilla houses one Rh50^+^ ORN [30]. As in ac1 sensilla, we detected Amt in both Rh50^+^ neurons and support cells in the sacculus (Figures 3B and 3C). Sensilla in sacculus chamber III house two neurons [30]. One neuron expresses olfactory receptor Ir64a and responds to acids [31], whereas markers for the second neuron have not been reported, precluding its functional analysis. We found that Ir64a expressing ORNs do not express Amt (Figure 3D), suggesting that sacculus III Amt/Rh50^+^ and Ir64a^+^ neurons are distinct. Consistent with this interpretation, previous reports indicate that Ir64a^+^ ORNs project to DC4 and DP1m antennal lobe glomeruli [31], whereas we find that Rh50^+^ ORNs project to VM6 (Figures 1J-1L). Chamber III sensilla are proposed to contain only one olfactory neuron because the dendrite of the second neuron does not fully extend into the sensillum lumen in electron microscopy images [30]. This raised the question whether the sacculus Amt/Rh50^+^ neurons are ammonia-sensitive like their ac1 counterparts. Sacculus sensilla are inaccessible for electrophysiological recordings, but their function can be assayed using calcium imaging. Our initial characterization of Amt/Rh50^+^ ORN activity was carried out with antennal lobe GCaMP imaging, but did not distinguish sacculus and ac1 neuron responses. We therefore turned to transcuticular imaging of *Rh50>GCaMP* fly antennae, where the two neuron populations can be segregated by location (Figures 3E and 3F). Similar to ac1 neurons, sacculus neurons showed dose-dependent responses to ammonia (Figures 3G-3J), providing evidence that the two neuron populations are both molecularly and functionally similar.

**Figure 3.**
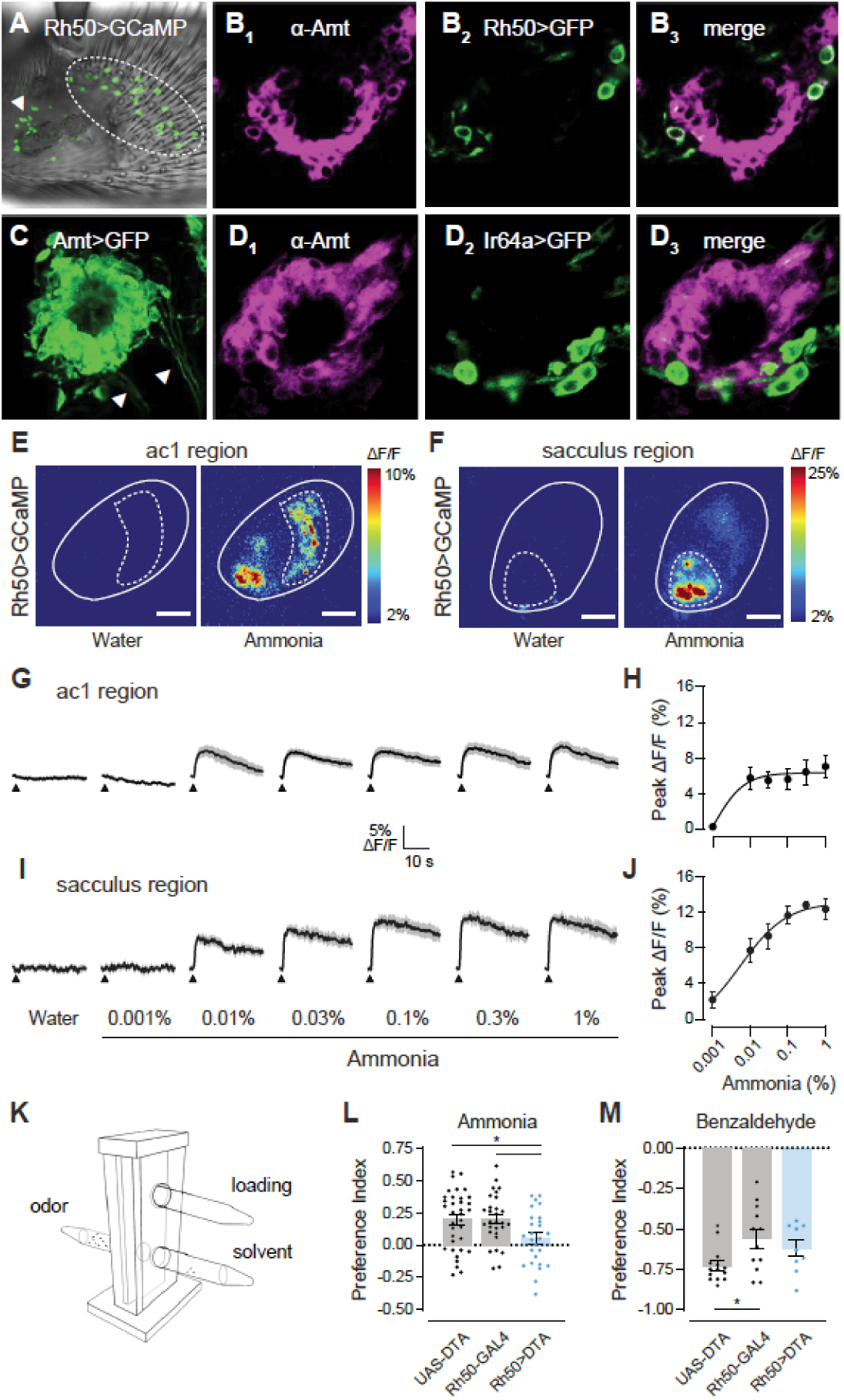
Two populations of Amt/Rh50^+^ ORNs mediate ammonia sensing. (A) Whole mount image of an antenna from a *Rh50>GCaMP7s* fly. Rh50^+^ neurons (green) are found on the ac1 region of the antennal surface (dotted line) and surrounding the sacculus (arrowhead). (B) Close-up view of sacculus chamber III in an antennal section from a *Rh50>GFP* fly stained with an anti-Amt (magenta, B_1_) and anti-GFP (green, B_2_). The merged image (B_3_) shows that Rh50^+^ ORNs co-express Amt. (C) High gain confocal image of sacculus chamber III in an antennal section from an *Amt>GFP* fly stained with an anti-GFP antibody. Labeled axons (arrowheads) emerge from clusters of GFP^+^ cells. (D) Immunostaining for Amt (magenta, D_1_) and GFP (green, D_2_) on antennal sections from *Ir64a>GFP* flies. (D_3_) Merged image showing that Ir64a^+^ ORNs do not co-express Amt. (E) Pseudocolored heat maps of calcium responses in the ac1 region (dotted line) of the antenna (solid outline) of *Rh50>GCaMP7s* flies to either water or 0.1% ammonia. (F) Similar to (E) but acquired at a different depth and location to focus on the sacculus region (dotted outline). (G) Traces of the mean calcium responses (black) ± SEM (gray) in the ac1 region. Arrowheads indicate time when the 250 ms odor stimulus was applied. (H) Dose-response curve of the peak ac1 calcium responses. (I) Calcium responses in the sacculus region. (J) Dose-response curve of the peak calcium responses in the sacculus (n= 6-7 flies for transcutical imaging). (K) T-maze assay schematic showing the elevator in the lower position with flies moving between the odor and solvent arms. The loading tube is above and is accessible with the elevator in the upper position. (L and M) Preference indices of *Rh50>DTA*, *UAS-DTA* and *Rh50-GAL4* flies when given the choice between 0.3% ammonia and water (L) or between 1% benzaldehyde and paraffin oil (M). Each dot represents one assay of ∼20-30 flies (n=26-35 assays for ammonia and 9-12 for benzaldehyde).

### Amt/Rh50^+^ ORNs contribute to NH_3_ attraction

Like many other insects, fruit flies are attracted to low levels of ammonia [12]. We examined the contribution of Amt/Rh50^+^ ORNs to ammonia seeking behavior using a T-maze two-choice assay in which naïve flies are given a short time to navigate towards or away from an odorant (Figure 3K). For these assays, we used flies in which Amt/Rh50^+^ ORNs were ablated by diphtheria toxin, the same genotype used for electrophysiological recordings (Figures 2C and 2D). Parental control lines (*UAS-DTA* and *Rh50-GAL4*) showed a preference for 0.3% ammonia, and this attraction was significantly reduced ∼75% in flies in which these transgenes were combined to eliminate Amt/Rh50^+^ ORNs (Figure 3L). We tested their general odor responsiveness and locomotor ability using benzaldehyde, an aversive odor. Benzaldehyde was highly repellent to both parental control lines, with their responses varying slightly (Figure 3M). Benzaldehyde aversion was unchanged in *Rh50>DTA* flies relative to parental controls (Figure 3M), confirming their overall odor sensing capabilities. Together, these data indicate the importance of Amt/Rh50^+^ ORNs in *Drosophila* ammonia attraction behavior.

### NH_3_ reception does not involve IRs/ORs

What is the ammonia receptor in these Amt/Rh50^+^ neurons? Nearly all insect olfactory neurons utilize a member of either the IR or OR receptor families [1, 5]. Individual tuning receptors that bind odorants each rely on a co-receptor for function, with odor responses absent in co-receptor mutants [2, 32, 33]. Members of the OR family rely on Orco, whereas IR receptors rely on Ir25a, Ir8a, or Ir76b [2, 32, 33]. We therefore asked whether any of these co-receptors are expressed in the ac1 Amt/Rh50^+^ ORNs. By using transgenic reporters for each of the co-receptors, we found that only Ir25a was co-expressed with Rh50 (Figures 4A-4D). However, single-sensillum recordings on ac1 sensilla revealed that ammonia-induced responses were unchanged in Ir25a mutant flies (Figure 4E), in agreement with a previous study [2]. Consistent with our expression data (Figures 4A-4D), ac1 ammonia responses were also not reduced in mutants for any of the three other co-receptors (data not shown).

**Figure 4.**
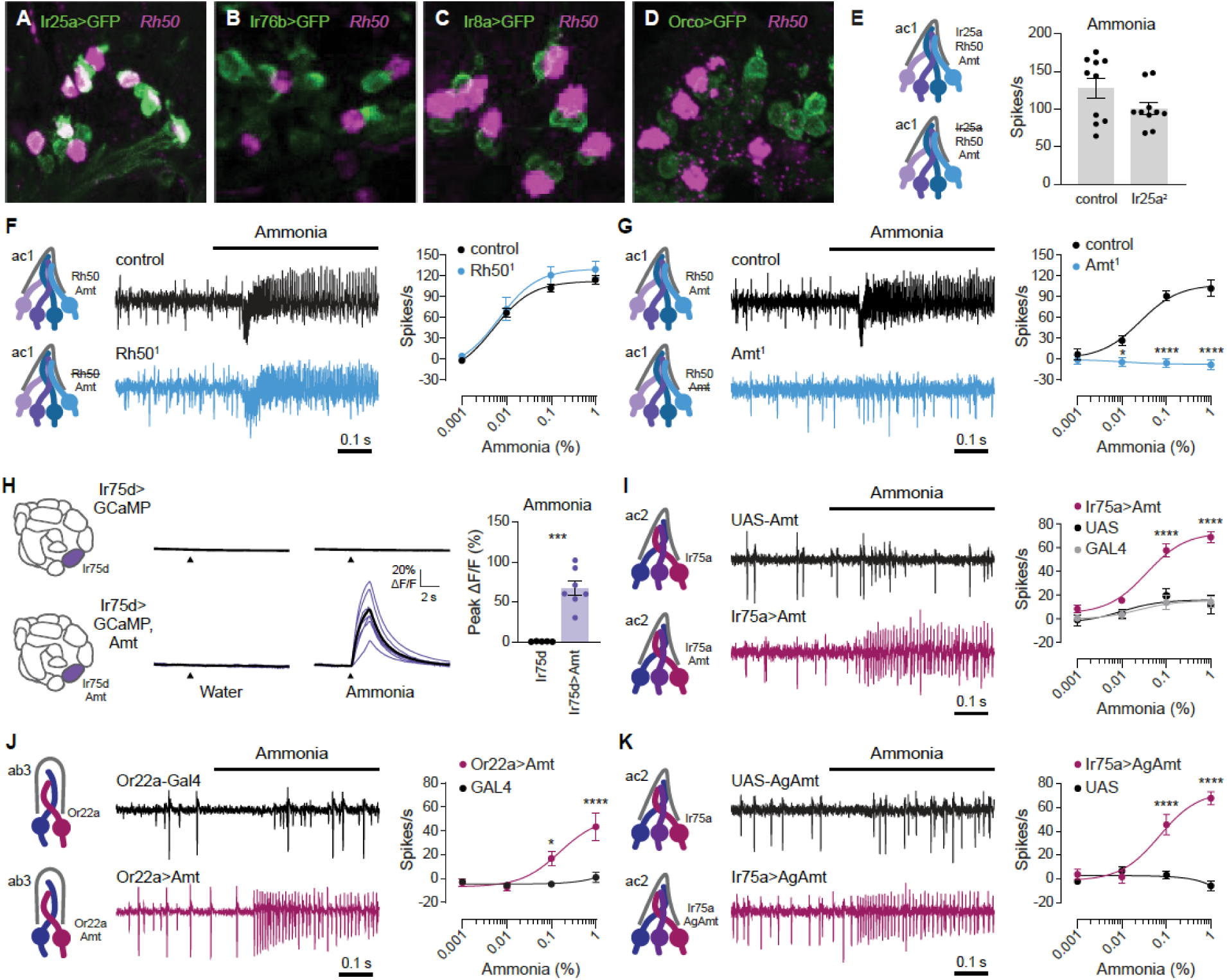
Amt transporter serves as an olfactory receptor for ammonia. (A-D) Confocal images of antennal sections labeled with an antisense probe for Rh50 (magenta) and an antibody against GFP (green) driven by *Ir25a-GAL4* (A), *Ir76b-GAL4* (B), *Ir8a-GAL4* (C), or *Orco-GAL4* (D). (E) Action potential responses to 0.1% ammonia in ac1 sensilla in control flies and those lacking Ir25a (n= 10 sensilla). (F) Action potential responses to 0.1% ammonia in ac1 sensilla in *Rh50^1^* mutants (blue) and control flies (black). Left, representative traces to 0.1% ammonia. Right, dose-response curve (n= 8-10 sensilla). (G) Action potential responses to ammonia in ac1 sensilla in Amt^1^ mutants (blue) and control flies (black). Left, sample traces to 0.1% ammonia. Right, dose-response curve (n= 8 sensilla). (H) Left, antennal lobe calcium responses to water and 0.01% ammonia in axon termini of *Ir75d>GCaMP6s* flies, with and without ectopic expression of Amt. Purple lines are responses in individual flies, and black lines are the mean response. Right, quantification of peak responses in *Ir75d>GCaMP6s* (black) and *Ir75d>GCaMP,Amt* (purple) flies (n= 5 and 7 flies). (I) Action potentials elicited by ammonia in ac2 sensilla in *Ir75a>Amt* flies (crimson) and control lines *Ir75a-GAL4* (grey) and *UAS-Amt* (black). Left, sample traces of 1% ammonia responses. Right, dose-response curve (n= 7-9 sensilla). (J) Action potentials elicited by ammonia in ab3 sensilla in *Or22a>Amt* flies (crimson), and *Or22a-GAL4* controls (black). Left, sample traces of 1% ammonia responses. Right, dose-response curve (n= 9-11 sensilla). (K) Spiking responses to ammonia in ac2 sensilla in *Ir75a>AgAmt* flies (crimson) and control *UAS-AgAmt* (black). Left, representative traces of responses to 1% ammonia. Right, dose-response curve (n= 7-8 sensilla).

### Amt acts as an ammonia receptor

The observation that none of the OR or IR co-receptors was required for ac1 ammonia responses raised the possibility that the Amt/Rh50^+^ ORNs utilize a non-canonical receptor to detect ammonia. We wondered whether the ammonium transporters might fulfill this role. Ammonium transporters in other species are highly selective for ammonium, with little to no transport of larger amines [34, 35], in accordance with the narrow tuning of Amt/Rh50^+^ ORNs to ammonia (Figure 2A). Further, ammonium transporters are electrogenic, mediating inward, depolarizing NH_4_^+^ influx [17, 18, 35, 36]. Given that ammonia (NH_3_) gas is expected to form ammonium ions (NH_4_^+^) in the sensillum lymph, these cations could be transported through either Amt or Rh50 and depolarize the ORNs. In this manner, the transporters could act as ammonia-selective receptors.

If this model is correct, ammonia responses should require the presence of either Amt or Rh50. To test the function of Rh50, we generated a Rh50 mutant which eliminated Rh50 expression (Figure S4). Spike responses to all concentrations of ammonia persisted in ac1 sensilla, if anything trending slightly higher than controls (Figure 4F). In contrast, *Amt^1^* mutants entirely lack ammonia responses at all concentrations tested (Figure 4G), as observed previously [14].

To determine whether Amt operates as an ammonia receptor, we tested whether ectopic Amt expression can confer ammonia sensitivity to ammonia-insensitive neurons. Indeed, expression of Amt in Ir75d^+^ neurons conferred robust antennal lobe calcium responses to ammonia that were absent in controls (Figure 4H). Further, ectopic expression of Amt in Ir75a^+^ ORNs, found in ac2 coeloconic sensilla, produced dose-dependent spiking responses to ammonia that were not observed in parental control lines (Figure 4I). Finally, Amt misexpression conferred a similar ammonia sensitivity to basiconic Or22a^+^ ORNs (Figure 4J), demonstrating that Amt function requires neither morphological nor molecular features specific to coeloconic sensilla. Together, these data indicate that Amt is an olfactory receptor for ammonia in *Drosophila*.

Amt is evolutionarily conserved, with orthologs found in multiple insect species, including the malaria vector *Anophele*s mosquitoes [16, 18]. Misexpression of AgAmt in Ir75a^+^ ORNs produced dose-dependent responses to ammonia (Figure 4K), similar to those induced by *Drosophila* Amt (Figure 4I). This raises the possibility that ammonium transporters also operate as ammonia receptors in other insect species.

## Discussion

Our identification of two previously unstudied populations of olfactory neurons expressing Amt and Rh50 transporters has paved the way to a new understanding of ammonia detection in *Drosophila*. Neuron ablation and calcium imaging experiments demonstrate that these ORNs mediate peripheral ammonia responses, and loss of these neurons greatly impairs ammonia seeking behavior. Unexpectedly, these neurons utilize the ammonium transporter Amt as a non-canonical odor receptor.

In insects, ORNs detect odors using IR and OR olfactory receptors, ligand-gated cation channels that are generally tuned to multiple structurally related odorants. Our loss and gain-of-function experiments reveal that the newly identified Amt/Rh50^+^ ac1 ORNs instead use an ammonium transporter as their receptor (Figures 4G-4K); the molecularly and functionally similar sacculus ORNs likely operate similarly. The narrowly tuned ammonia response in these ORNs (Figures 2A and 2D) is consistent with the strict selectivity of ammonium transporters [34, 35]. Amts also distinguish NH_4_^+^ ions from similarly sized K^+^ ions [18, 35, 36], an advantage considering the presence of >100 mM K^+^ in the sensillum lymph bathing ORN dendrites. We propose that a simple electrogenic influx of ammonium ions leads to ORN depolarization, but we cannot rule out an ammonium-initiated signaling cascade analogous to the mechanism for sour sensing in vertebrate taste cells, in which H^+^ influx through Otop1 proton channels closes acid-sensitive K^+^ channels [37]. However, an ammonium signaling cascade is unlikely because it would require the components to be widely expressed in molecularly diverse ammonia-insensitive neurons in both basiconic and coeloconic sensilla.

The ammonia-sensing ORNs in *Drosophila* express both Amt and Rh50, members of the two main branches of the ammonium transporter family. Ammonium transporters in plants, bacteria, and fungi serve to uptake ammonium for biosynthesis of nitrogenous molecules, whereas animal Rh proteins play a role in ammonia excretion and acid-base homeostasis [16, 17]. Our findings indicate a novel role for Amt ammonium transporters as chemosensory receptors in animals. In contrast, we did not find a significant role for Rh50 in ammonia sensing, perhaps due to its >20-fold lower expression in antennae than Amt [14]. Rh50 may also be unable to function as an ammonia receptor because most Amts transport ammonium at low micromolar concentrations whereas Rh proteins require millimolar levels [38]. The role of Amt proteins in support cells is likewise unclear; Amt misexpression into neurons in coeloconic and basiconic sensilla that lack Amt in support cells is sufficient to induce ammonia-sensitivity (Figures 4I and 4J). Support cell Amt may affect the magnitude of the ORN response, a possibility suggested by the somewhat smaller ammonia responses in ac2 versus ac1 ORNs (Figures 4G and 4I).

Ammonia-sensitive ORNs are found in every insect species examined, and many insects are attracted to low levels of ammonia. Several lines of evidence suggest that Amt orthologs are widely used by insects as ammonia receptors, and therefore likely mediate this behavior. First, Amt proteins are highly conserved, with for example, *Drosophila* and *Anopheles* Amts sharing 83% amino acid identity in the transmembrane regions. Accordingly, expression of *Anopheles* AgAmt conferred ammonia sensitivity to ORNs (Figure 4K), and a recent study detected Amt^+^ neurons in ammonia-sensitive olfactory sensilla in *Anopheles* mosquitos [19]. Additionally, Amt orthologs can be found in antennal transcriptomes from multiple, distantly related insect species [14]. Together these similarities suggest Amt transporters have taken on a new function as ammonia receptors in insects.

## Acknowledgments

We thank Tzumin Lee, Greg Suh, John Carlson, Richard Benton and the Bloomington *Drosophila* Stock Center (NIH P40OD018537) for fly lines. We thank the Developmental Studies Hybridoma Bank, NIH NICHD at the University of Iowa, for antibodies. The pBGRY plasmid was a gift from John Carlson and Tong-Wey Koh. The pU6-BbsI-chiRNA plasmid was a gift from Melissa Harrison & Kate O’Connor-Giles & Jill Wildonger. The pHD-DsRed plasmid was a gift from Kate O’Connor-Giles. We thank Yinan Xuan for assistance with transcutical imaging data analysis, Tiffany Liang for help with initial behavioral experiments, and Pratyajit Mohapatra for genomic sequencing validation of mutants. We thank Anastasios Tzingounis for discussions and comments on the manuscript. This research in K.M.’s laboratory was supported by NIH awards R21DC017868, R03DC015629, and R35GM133209. Research in C-Y.S.’s lab was supported by R01DC016466, R01DC015519 and R21DC108912. V.G. and S.S. were supported by the Max Planck Society. This research in J.M.J.’s laboratory was supported by NIH awards R01DC018570, R01NS116584, the Richard and Susan Smith Family Award for Excellence in Biomedical Research, the Klingenstein-Simons Fellowship Award in Neuroscience, and The Kavli Institute at Yale University. H.S.K. is supported by an NSF Graduate Research Fellowship.

## Author Contributions

K.M. conceived the project, with A.V. and C.Y.S. providing conceptual contributions. All authors contributed to experimental design, analysis, and interpretation of results. A.V. generated transgenic *Drosophila* lines and performed electrophysiological recordings. A.V., S.B., and V.G carried out histological studies. A.V. and S.B. performed behavioral experiments. H.K. and S.W. performed GCaMP imaging. S.B. carried out qPCR analysis. C.N.V. analyzed SBEM data. A.V and K.M. wrote the paper with input from all other authors.

## STAR Methods

### Lead Contact

Further information and requests for resources and reagents should be directed to and will be fulfilled by the Lead Contact, Karen Menuz.

### Materials Availability

All novel biological materials, including transgenic *Drosophila* strains, anti-Amt antibody, and plasmids, are available upon request.

### Data and Code Availability

All custom codes used for analysis are available upon request.

### Experimental Model and Subject Details

*Drosophila melanogaster* were reared on a standard cornmeal food at 25°C in a humidified incubator with a 12:12 hour light/dark cycle. The genotypes of fly lines used in this study are listed in Table S1. The *Ir92a^1^*, *Rh50^1^*, and *Ir25a^2^* mutant lines were outcrossed for at least five generations to a *Canton-S* background prior to electrophysiological recordings, as were transgenes in flies used for recordings in Figure 2 and behavior in Figure 3. The *Amt^1^* transposon mutation had been previously outcrossed to an isogenic *w^1118^* line [14, 40]. Experimental flies were between 2-12 days old. The genders and specific age ranges for different experiments are provided in the Method Details below.

### Method Details

#### Reporter line generation

Transgenic GAL4 reporter flies were generated using standard methods as described [14]. The 5’ and 3’ regions flanking the genes were amplified from bacterial artificial chromosomes (BACs) corresponding to the reference genome of *Drosophila melanogaster* [41]. MultiSite Gateway Pro recombination (Thermo Fisher) was used to assemble the 5’ and 3’ genomic regions with GAL4 in the pBGRY destination vector [42]. For *Rh50-GAL4*, the 5’ region included chromosome 3L: 4,907,401 to 4,910,693 and the 3’ region 3L: 4,918,898 to 4,924,642. For *Ir25a-GAL4*, the 5’ region included chromosome 2L: 4,835,726 to 4,834,655 and the 3’ region 3L: 4,830,990 to 4,827,634. For *Ir75a-GAL4*, the 5’ region included chromosome 3L: 17,829,014 to 17,817,922 and the 3’ region 3L: 17,815,667 to 17,811,236. PhiC31 integration (Best Gene, Inc.) was used to integrate the assembled GAL4 vectors into the *Drosophila melanogaster* genome. Two transgenic strains were made for each GAL4 in which the construct was incorporated into the second (attP40 landing site [43]) and third (attP2 landing site [44]) chromosomes.

#### Generation of Ir92a and Rh50 mutants

The *Ir92a^1^* and *Rh50^1^* mutant alleles were generated with CRISPR/Cas9 engineering and homology directed repair, similar to described previously [45]. The *Ir92a^1^* mutation deletes the terminal 323 amino acids of Ir92a (52% of the coding sequence), including all transmembrane domains. The *Rh50^1^* mutation eliminates 143 amino acids (29% of the coding sequence), including several transmembrane domains.

Guide RNA sequences were cloned into pU6-BbsI-chiRNA [46], using the Q5 Site-Directed mutagenesis Kit (New England Biolabs). Two gRNAs were used for targeting Ir92a (GGTCACCGAAGAACGGGCTA and GGACGCATCTCCCCGTGAAA) and one for Rh50 (GATCCAGTCTGTCCAGGTTC). A donor plasmid for homology directed repair for Ir92a was generated by cloning homology arms from wCS genomic DNA and inserting them into the pHD-DsRed-attP vector [47] to flank the 3xP3-DsRed marker. Rh50 homology arms were cloned from BAC R17L24 and inserted into a modified pHD-DsRed attP vector that contained a LexA reporter sequence 5’ to the 3xP3-DsRed site. For each gene, gRNA and donor plasmids were injected into embryos of P{nos-Cas9.R}attP40/ CyO flies [48] (Best Gene, Inc.).

Flies in which the donor plasmid had integrated into the genome were identified by ocular DsRed expression. We used PCR and sequencing to validate the location of donor plasmid integration. The Ir92a donor plasmid integrated as expected, with the 3xP3-DsRed marker replacing genomic region 3R: 20,342,366-20,344,610. The Rh50 donor plasmid did not integrate as designed. Instead genomic region 3R: 4,913,350-4,914,409 was replaced by sequences from the donor plasmid extending from the ampicillin resistance region through the LexA and 3xP3-DSRed regions. Flies were genotyped (Figures S3 and S4) using the following primers:

Ir92a pair 1: TGTATGGCCGGTAGGATCTC and ACCTCCTTGATCGAAACCCT

Ir92a pair 2: GGCAAGAATGCGAACAAAT and TGGTTTGTCCAAACTCATCAA

Rh50 pair 1: CCTCTCCCTGGAGAACATCA and CCCTCTAGCTTTCCCGTTTC

Rh50 pair 2: CTGTTCATGGCTGCTCTAGT and CTGAGATAGGTGCCTCACTG

#### Quantitative reverse transcriptase-PCR (qRT-PCR)

Heads from ten female Rh50^1^ and wCS flies aged 6-7 days were dissected over liquid nitrogen for RNA extraction. The heads were crushed with a pestle and passed through a QIAshredder column (Qiagen), and then total RNA was extracted with the RNeasy Micro Kit (Qiagen). 100 ng of RNA was reverse transcribed into cDNA using the iScript gDNA Clear cDNA Synthesis Kit (Bio-Rad). In parallel, RNA was separately processed without the reverse transcriptase to control for any gDNA contamination. qRT-PCR reactions were run in triplicate on a CFX96 thermocycler (Bio-Rad) with SsoFast EvaGreen Supermix (Bio-Rad) containing cDNA from 5 ng of RNA and Rh50 primers AATGAGCAGTGTGACAGCGA and CATTGCCTCCGCCATTTACG. Expression in each sample was normalized to a housekeeping gene, eIF1A, detected with primers ATCAGCTCCGAGGATGACGC and GCCGAGACAGACGTTCCAGA.

#### FISH and antennal immunocytochemistry

Male and female flies 7-10 days old were placed in an alignment collar. Their heads were encased with OCT (Tissue-Tek) in a silicone mold, frozen on dry ice, and snapped off. These head blocks were stored at −80°C. A cryostat was used to collect 20 μm sections.

Immunocytochemical staining was carried out identically as done previously for labellar sections [45]. Transgenic GFP expression was detected with mouse anti-GFP antibody (1:500, Roche) and donkey anti-mouse Alexa Fluor 488 (1:500, Thermo Fisher). A rat anti-elav antibody (1:10, Developmental Studies Hybridoma Bank) was visualized with goat anti-rat Alexa Fluor 568 (1:500, Thermo Fisher). A guinea pig anti-Amt antibody [45] (1:200) was detected with goat anti-guinea pig Alexa Fluor 568 (1:500, Thermo Fisher). Combined FISH and immunocytochemistry staining were carried out as described previously [14]. The digoxigenin (DIG)-labelled Rh50 probe was generated from a plasmid containing the full-length cDNA sequence corresponding to Rh50-RA isoform (Drosophila Genomics Resource Center) [49]. This plasmid was digested with XhoI for T7 transcription (sense probe) and EcoRV for SP6 transcription (anti-sense probe) for labelling with the DIG RNA Labeling Kit (SP6/T7) (Roche). Stained sections were imaged on a Nikon A1R confocal microscope in the UConn Advanced Light Microscopy Facility. Stacks of images (0.5 µm z-step size) were collected and analyzed with ImageJ/Fiji software [50].

#### Antennal lobe immunocytochemistry and 3D reconstruction

Brains of 7 day old female and male flies were dissected in PBS and directly transferred to 4% PFA with 0.1% Triton-X (PBST) for fixation over 2 hours on ice. After three 15 minute washes with PBST and blocking in 5% normal goat serum (NGS) in PBST the primary antibodies mouse anti-brp (1:30, Developmental Studies Hybridoma Bank) and rabbit anti-GFP (1:500, Thermo Fisher) were applied in PBST with 5% NGS for 4 days nutating at 4°C. Next, the brains were washed four times for 15 minutes with PBST, blocked again with 5% NGS in PBST and incubated with the secondary antibodies goat anti-mouse Alexa Fluor 633 and goat anti-rabbit Alexa Fluor 488 (1:250 each, Thermo Fisher) in PBST with 5% NGS for 5 days nutating at 4°C. After four final washing steps with PBST, brains were mounted in VectaShield (Vector Labs). Stacks of immunostained brains were scanned on a Zeiss cLSM 880 confocal microscope with a z-step size of 0.44 µm. The confocal stacks were manually reconstructed as label fields in AMIRA (6.7, Thermo Fisher) and identified on the basis of the published *Drosophila melanogaster* atlases [51]. Surface renders were created and smoothed in AMIRA with the surface gen and smooth surface tools.

#### *In vivo* microscopy of antennal lobes

To visualize the Rh50 positive ORNs *in vivo*, an open head dissection was utilized. The flies were anesthetized on ice and then glued to a plastic holder to reduce movement. A wire was installed to bend the antennae forward and maximize the cutting area at the vertex. After adding a plastic coverslip with a hole, and sealing this hole around the vertex with two-component silicone, the vertex was cut open under saline. The trachea and fat tissue inside the head were removed to reveal a clear view onto the central brain. The image shown in Figure 1K was obtained with a Zeiss cLSM 710 multi photon microscope.

#### Serial block-face scanning electron microscopy (SBEM) analysis

Coeloconic sensilla were identified in SBEM datasets that were previously generated [21, 22], based on their signature cuticular finger structure [20]. The number of ORNs within each coeloconic sensillum was determined by examining the EM images using IMOD v4.9 [52]. Segmentation was performed manually. The 3D models were then generated using the IMOD command “imodmesh” and smoothed using the “smoothsurf” command. The sensillum shown in Figure 1I was identified in the Or88a-labeled dataset [21].

#### Whole-mount Confocal Imaging

7-day old female flies were anesthetized on ice before their antennae were removed for fixation with 4% paraformaldehyde/PBS (MPX00553, Fisher Scientific) for 20 min at room temperature. After washing three times with 0.3% PBT (PBS with 0.3% Triton X-100), the samples were mounted in FocusClear (CelExplorer Labs Co.) and imaged on a Zeiss LSM 880 confocal microscope using 40x, N.A.1.2 C-Apochromat water-immersion objective lens. Airyscan images were processed with ZEN (Zeiss) and the brightness and contrast were further adjusted using ImageJ/Fiji.

#### Behavioral assay

T-maze behavioral assays were performed using custom-built acrylic apparatuses. At 0-4 days post-eclosion, flies were sorted into groups of 16 males and 16 females of the appropriate genotype. The vial of flies was tested at 7-12 days post-eclosion. Flies were starved in empty vials with 2 mL of distilled water to moisten the foam plugs ∼22 hours prior to the assay. Assays were conducted within a three hour time window to control for any circadian effects. Two hours prior to the start of the assay, fly vials were moved to a darkened quiet room to acclimate to the room temperature and light level. Antibiotic assay discs (13 mm, Whatman) were placed tightly into the bottom of 15-mL conical tubes (DOT Scientific) using a metal spatula, and 50 µl of either odorant or solvent were pipetted onto the disc. The tubes were then capped and the odorant allowed to volatilize for 30 minutes before a set of assays began.

A set of ∼10 consecutive assays was carried out over a period of ∼45 minutes in a dark room lit with dim red light. The assay chambers were placed in a cardboard box to further limit visual cues. For each assay, ∼20-30 flies were transferred with a funnel from a vial into a 15-ml centrifuge tube, which was then screwed on the upper opening of the T-maze central tower. Flies were then tapped into the elevator, which was at its topmost position to align with the loading tube. The elevator was partially lowered, and flies were given one minute to acclimate. Towards the end of this time, odorant and solvent tube “arms” were screwed into the T-maze, with the positions of the odorant and solvent arms alternating between assays. The elevator was then lowered further so that flies had access to both the odorant and solvent arms. Flies had one minute to chemotax into the arms before the elevator was raised to trap flies in the arms and prevent further movement. Preference index was calculated as the number of flies entering the solvent arm subtracted from the number of flies entering the odorant arm, and this value divided by the total number of flies in the assay including those that remained in the elevator hole. Each set of flies and conical tubes was only used for one assay. Each set of assays tested flies of multiple genotypes in a random order.

#### Odorants

Odorants were ammonium hydroxide (Fisher or VWR International), trimethylamine (Sigma), dimethylamine (Sigma), ethylamine (Sigma), butylamine (Sigma), cadaverine (Sigma), spermidine (ACROS), isoamylamine (ACROS), and benzaldehyde (Sigma). All odorants were diluted to generate 10% stocks, and serial dilutions were used to generate lower concentrations. All odorants were diluted in water, except benzaldehyde, which was diluted in paraffin oil (ACROS). Ammonia was used at concentrations from 0.0001% to 1% for electrophysiology and transcuticular imaging. Other odorants used for electrophysiological experiments were at 1% concentration. All odorants were used at a 0.01% concentration for antennal lobe GCaMP imaging. For behavioral assays, ammonia was used at 0.3% and benzaldehyde at 1%.

#### Electrophysiology

Single-sensillum electrophysiological recordings were generally performed on 3-5 day old female flies as described [53, 54]. Flies for diphtheria-toxin ablation experiments were aged 8-11 days to ensure complete neuron ablation. In brief, flies were wedged in the narrow tip of a 200 µl pipette tip, exposing the antennae and a portion of their head. One antenna was stabilized between a tapered glass electrode and a coverslip. The prep was placed on a BX51WI microscope (Olympus) under a continuous 2,000 mL/min stream of humidified, purified air. A borosilicate glass electrode filled with sensillum recording solution [55] was placed into the eye as a reference electrode, and an aluminosilicate electrode filled with the same solution was inserted into individual sensilla. Sensilla classes were identified based on their known location on the antenna and their response profile to a small number of diagnostic odorants. Up to four sensilla were recorded per fly. Extracellular action potential recordings were collected with an EXT-02F amplifier (NPI) with a custom 10x gain headstage. Data were acquired and AC filtered (300-1,700 Hz) at 10 kHz with a PowerLab 4/35 digitizer and LabChart Pro v8 software (ADInstruments).

Odorant cartridges were prepared by placing a 13 mm antibiotic assay disc (Whatman) into a Pasteur pipette, pipetting 50 µl of odorant solution onto the disc, and enclosing the end with a 1 mL pipette tip. Cartridges were allowed to equilibrate for at least 20 minutes. Cartridges were used no more than four times, with at least 10 minutes recovery between re-use for trials on different sensilla. Odorants were applied for 500 ms at 500 mL/min after inserting the cartridge into a hole in the main airflow tube. Odor delivery was controlled by LabChart, which directed the opening of a Lee valve (02-21-08i) linked to a ValveBank 4 controller (Automate Scientific). A ten second recording was collected, with a one second baseline period before odor application. Each sensillum was tested with multiple odorants, with at least a 10 second rest period between odor applications.

Action potentials were detected offline using LabChart Spike Histogram software. Spikes were sorted by their amplitude in basiconic recordings, whereas all spikes from the 3-4 ORNs were summed in coeloconic recordings due to their similar sizes. Action potentials were counted over a 500 ms window, 100 ms after stimulus onset due to the line delay for the odor to reach the antenna. Solvent corrected odor responses were calculated as the number of spikes induced by the odor after subtracting the number of spikes produced by stimulating with water alone.

#### Calcium imaging of ORN axon terminals in the antennal lobe

Genetically encoded calcium indicators belonging to the GCaMP family were used to visualize the odor-evoked calcium activity in the ORN axon terminals of interest in the antennal lobe. All experiments were conducted in virgin female flies aged 2-11 days post-eclosion. Experimental flies were collected shortly after eclosion and housed in small groups at 25°C under a 12 hour light and 12 hour dark cycle until imaging.

Flies were prepared for imaging as previously described [56]. The external saline contained 103 mM NaCl, 3 mM KCl, 5 mM TES, 8 mM trehalose ·2H2O, 10 mM glucose, 26 mM NaHCO3, 1 mM NaHPO4 ·H2O, 4 mM MgCl2 ·6H2O, and 1.5 mM CaCl2 ·2H2O and was adjusted to a pH of 7.25 and osmolarity of ∼270 mOsm. The fly was first head-fixed into a small hole in a thin sheet of stainless steel foil, such that its antennae protruded beneath the foil (to stay dry) while the rest of the head was submerged in the external saline solution above the foil. To target Rh50 ORN axons (Figure 2A), a small patch of cuticle dorsal to the brain was removed and fat, air sacs, and trachea were removed to obtain good optical access. To target Ir92a (Figure 2B) and Ir75d (Figure 4H) ORN axons the head was rotated 180° such that the proboscis pointed up. The proboscis was then removed, along with fat, air sacs, and trachea. In additional experiments, we imaged from Rh50 ORN axons ventrally and found no qualitative difference between the resulting response traces and those obtained dorsally (data not shown).

Following the dissection, the metal foil containing the head fixed fly and the external saline was mounted on an epifluorescence microscope under a 40x water-immersion objective lens. The fly was positioned under the microscope such that its antennae faced a constant stream of charcoal-filtered carrier air, delivered at the rate of ∼1360 mL/min. Odor stimuli were prepared by diluting stock chemicals in distilled water in 2mL vials. To deliver the odor stimulus into the carrier stream, a computer-controlled valve diverted a small amount of the carrier stream into the headspace of the odor vial, at the rate of ∼6.2 mL/min, before rejoining the carrier stream. GCaMP was excited with a 470 nm LED at 5% power, corresponding to 0.367 mW at the sample (CoolLED pE-100). The emitted fluorescence was collected by a Hamamatsu digital camera (model C1140-42U30) using HCImageLive software. Each imaging trial lasted 15 seconds at the frame rate of 16.67 Hz, with the odor delivery starting at 4 seconds into a given trial and lasting for 2 seconds.

Individual CXD image files were converted to TIF files using the batch process function from ImageJ software. TIF-converted data for each fly were concatenated into an image stack for each stimulus condition using a custom MATLAB script. For each stimulus condition, the ROI was defined by visually inspecting the image stack and tracing around the perimeter of the ORN axon terminal. The same ROI was applied to every frame of each stack. ΔF/F was calculated as (F_signal_ − F_0_) / F_0_, where F_signal_ is the instantaneous fluorescence pixel value averaged over the entire ROI and F_0_ is the averaged pixel value in the same ROI over the 1 second preceding the stimulus onset. In each fly, three trials were conducted for each stimulus condition; ΔF/F was averaged over each trial. These ΔF/F traces were then averaged across flies.

#### Transcuticular antennal calcium imaging

Female flies aged 7 days were used for antennal calcium imaging experiments. To prepare an antenna for recording, a fly was wedged into the narrow end of a truncated 200 µl pipette tip to expose the antenna, which was subsequently stabilized between a tapered glass microcapillary tube and a coverslip covered with double-sided tape. Images of different populations of Rh50-expressing neurons were acquired at different depths to focus on either the ac1 region or the sacculus region.

Images were acquired via Micro-Manager 1.4 (The Open Source Microscopy Software) with a CMOS camera (Prime 95B, Photometrics) and an upright microscope (Olympus, BX51WI) with a 50x air objective (NA 0.50, LMPlanFl, Olympus). Blue LED (470 nm, Universal LED Illumination System, CooLed pE-4000) was used to excite GCaMP7s. Image acquisition was at 10 Hz for ∼38 s. Light pulses (25-ms on, 75-ms off) were used to minimize photobleaching. Odorants (100 μl applied to a filter disc) were delivered from a Pasteur pipette via a pulse of air (200 mL/min) into the main air stream (2000 mL/min). A 250 ms pulse of odorant was applied 7 seconds after acquisition onset, with an interstimulus time interval of 2–3 min between each application.

All images acquired from the same antenna were first concatenated. A MATLAB function, NoRMCorre [57], was used for motion correction. For each antenna, ROIs were determined via a custom Python script based on the calcium responses to 0.3% ammonia. Briefly, the frame with the highest summed pixel value within the field of view was first generated (peak frame). A delta frame was then determined by subtracting the peak frame with the averaged pre-stimulus frame (2 seconds prior to odor stimulus). The delta frame was processed with a Gaussian filter to smooth the image. ROIs were then identified by applying a threshold (>70% of the highest pixel value) to the smoothed delta frame. The same ROIs were applied to all images acquired from the same antenna across different ammonia concentrations.

ΔF/F was calculated as (F_signal_ − F_0_) / F_0_, where F_signal_ is the instantaneous fluorescence pixel value averaged over the entire ROI and F_0_ is the averaged pixel value in the same ROI over the 2 second pre-stimulus period. In order to remove imaging noise, baseline ΔF/F—defined as the fluorescence level which is less than 10% of the highest pixel value of the entire image—was further subtracted from the ΔF/F of the ROIs. For each recording, the representative ΔF/F was determined by averaging the traces from three ROIs which had the highest ΔF/F values upon odor stimulation.

#### Quantification and Statistical Analysis

Data were analyzed using GraphPad Prism 8. Bar graphs depict the mean ± SEM overlaid with the individual data points. Dose response curves show the mean ± SEM and the curve fit to the Hill equation. Unpaired two-tailed t-tests were used to compare two genotypes, and one-way ANOVAs followed by Tukey post-hoc tests for three or more genotypes. Datasets involving multiple genotypes and multiple odorants or odorant concentrations were analyzed with two-way ANOVAs followed by Holm-Sidak post-hoc tests. Statistical details can be found in the figure legends. Statistical significance is presented as *p<0.05, **p<0.01, ***p<0.001, ****p<0.0001. Other comparisons were not significant (p>0.05).

---

**Figure S1.**
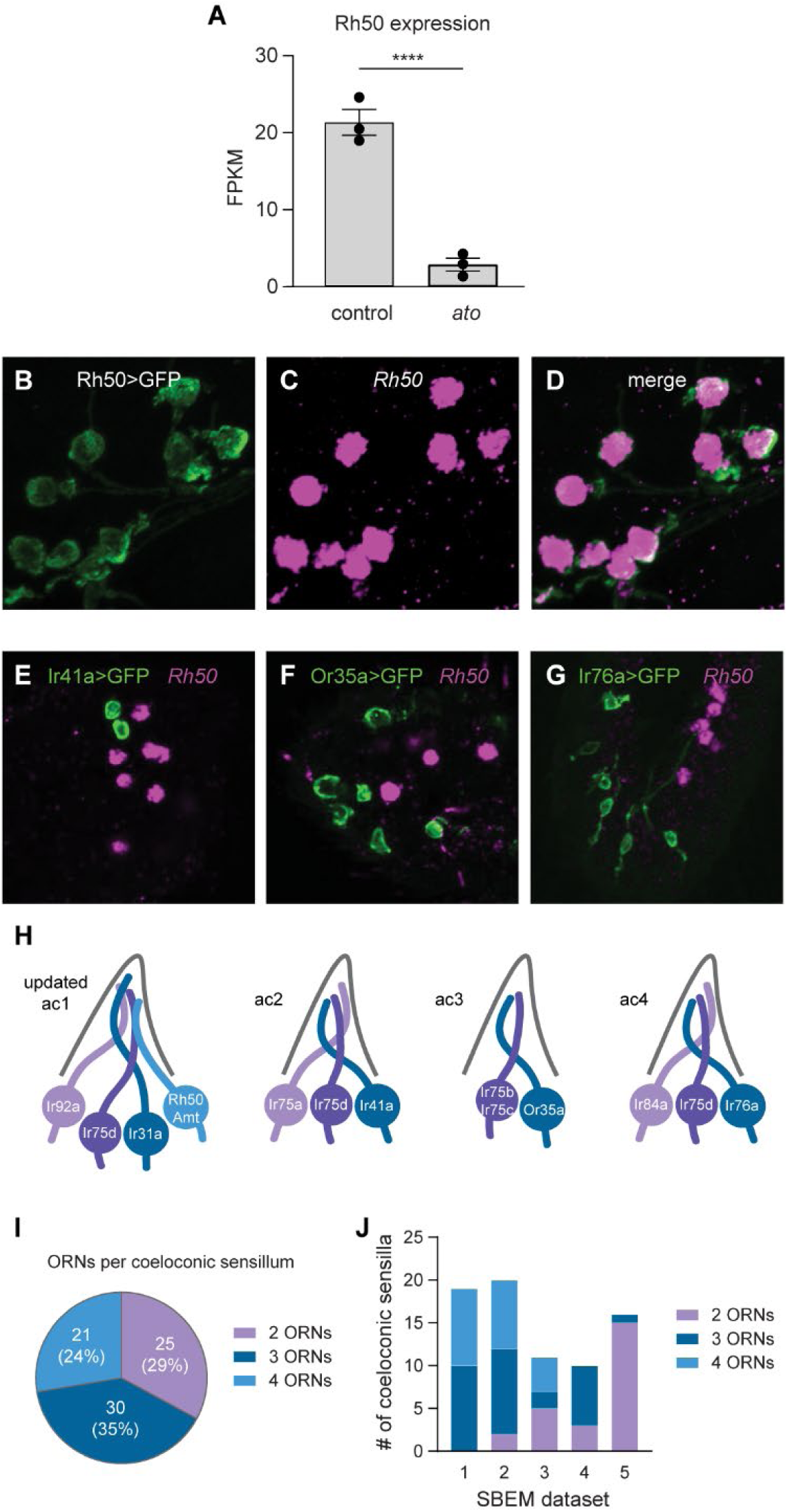
Antennal Rh50 expression and coeloconic sensilla organization (related to Figure 1). (A) Previous RNASeq analysis of Rh50 expression in antennal transcriptomes from control and *atonal* flies, which lack coeloconic sensilla (n=3 sets of ∼600 antennae) [14]. (B-D) Antennal section from a *Rh50>GFP* fly stained with an antibody against GFP (green, B) and an *in situ* hybridization probe to Rh50 (magenta, C). (D) Merged image shows overlapping labels. (E-G) Antennal sections from *Ir41a>GFP* (E), *Or35a>GFP* (F), and *Ir76a>GFP* (G) flies, which mark the location of ac2, ac3, and ac4 sensilla, respectively (H). The sections were labeled with an antisense probe for Rh50 (magenta) and an antibody against GFP (green). Rh50^+^ ORNs only rarely neighbor the GFP^+^ ORNs (green) labeled in these fly lines. (H) An updated model of the ORNs found in each coeloconic sensillum. (I) Number and percentage of coeloconic sensilla with two, three, or four ORNs found in five serial block face electron microscopy (SBEM) datasets. (J) Number of coeloconic sensilla with two, three or four ORNs found in each of the five SBEM datasets.

**Figure S2.**
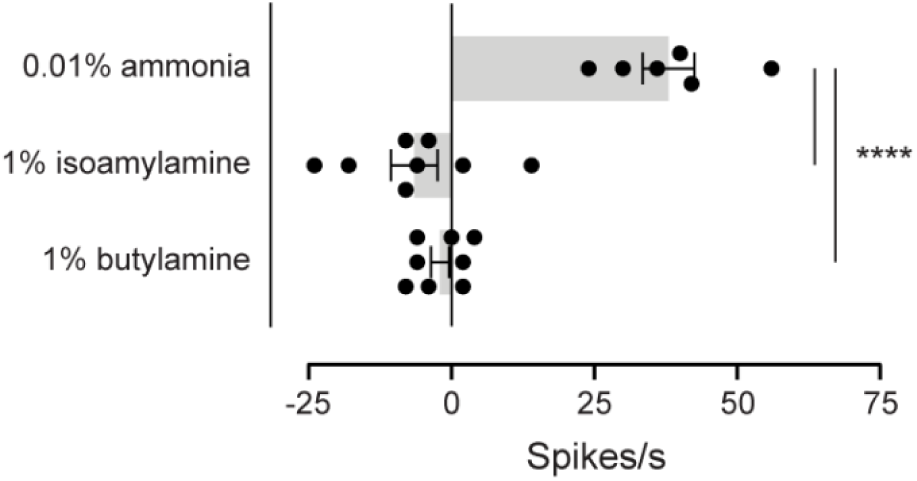
Analysis of ac1 pH sensitivity (related to Figure 2). Single sensillum recordings of action potentials in ac1 sensilla from a wild-type fly in response to ammonia (pKa 9.4) and higher concentrations of two more basic odorants: isoamylamine (pKa 10.6) and butylamine (pKa 10.8) (n=6-8 sensilla).

**Figure S3.**
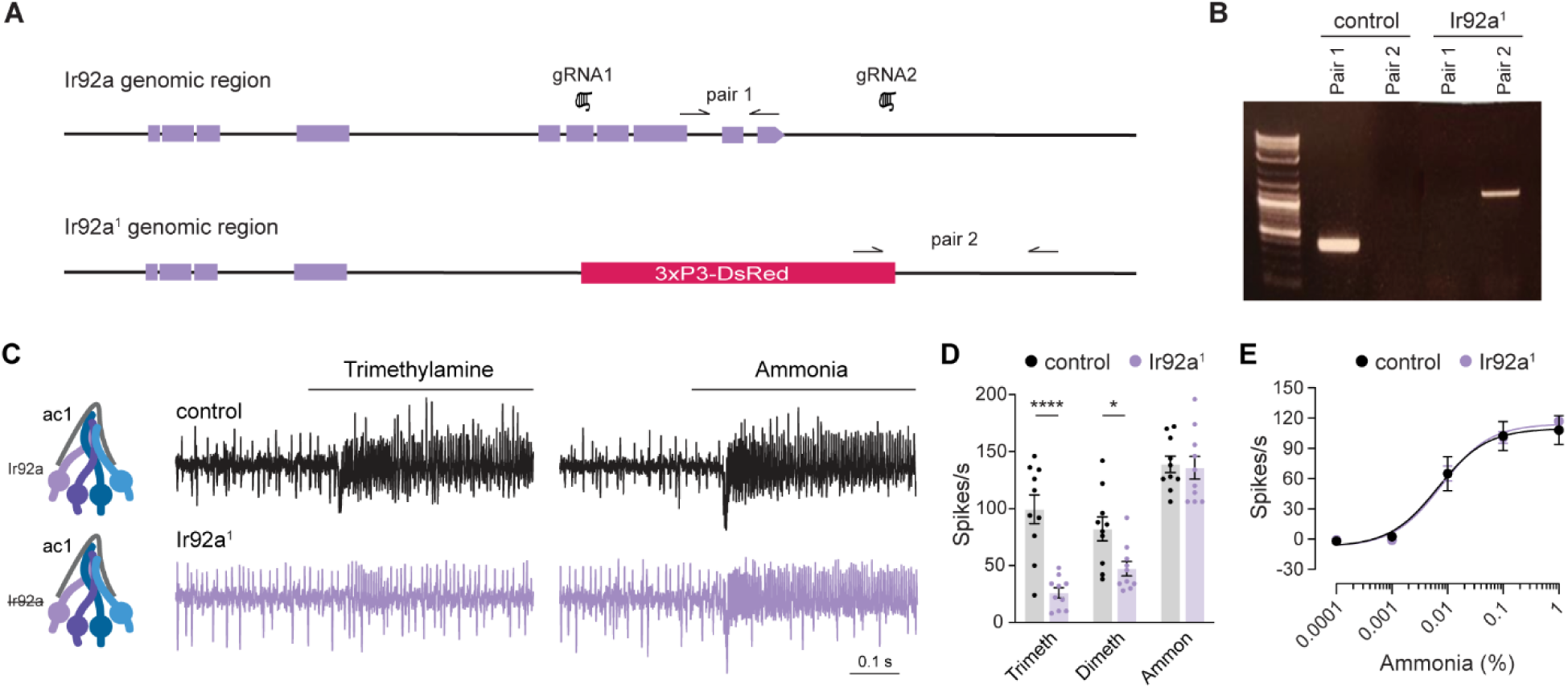
Generation and physiology of *Ir92a^1^* mutants (related to Figure 2). (A) Upper, a depiction of the Ir92a genomic region, with Ir92a exons in purple. The location of gRNAs 1 and 2 is shown. Lower, the Ir92a genomic region in *Ir92a^1^* mutant flies after homology directed repair. The region between gRNAs 1 and 2 was replaced with the 3xP3-DsRed marker. Genotyping primer pairs 1 and 2 are shown. (B) Agarose gel showing genotyping PCR bands from control and *Ir92a^1^* mutant flies with primer pairs 1 and 2. (C) Representative traces of extracellular recordings of action potentials elicited by 1% trimethylamine and 0.1% ammonia in ac1 sensilla from control flies (black) and *Ir92a^1^* receptor mutants (purple). (D) Quantification of solvent corrected odor responses in control (black) and *Ir92a^1^* (purple) flies (n= 10 sensilla). (E) Dose-dependent responses to ammonia in control (black) and *Ir92a^1^* flies (purple) (n= 8-10 sensilla).

**Figure S4.**
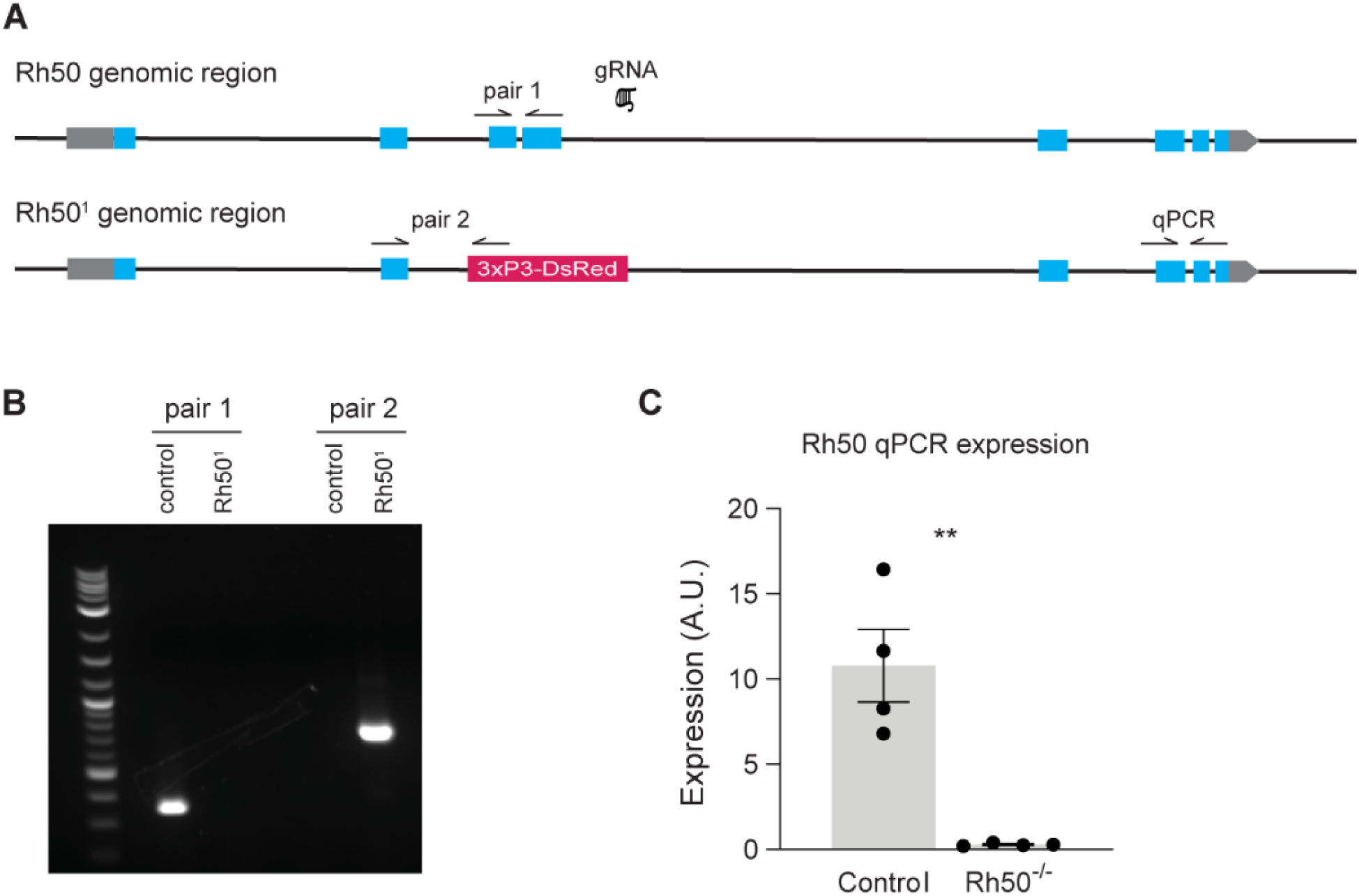
Generation and validation of *Rh50^1^* mutants (related to Figure 4). (A) Upper, a depiction of the Rh50 genomic region, with Rh50 exons in blue. The location of the gRNA is shown. Lower, the location of the 3xP3-DsRed insertion in the Rh50 genomic region in *Rh50^1^* mutant flies as determined with PCR and sequencing. Exons three and four are eliminated. Genotyping primer pairs 1 and 2 and qPCR primers are also shown. (B) Agarose gel showing genotyping PCR bands from control and *Rh50^1^* mutant flies with primer pairs 1 and 2. (C) qPCR analysis of Rh50 expression in cDNA from heads of control and *Rh50^1^* flies (n=4 biological replicates).

**Table S1.** *Drosophila* genotypes used in this study.

| ***Drosophila* stock name** | **Figure** |
| --- | --- |
| *w; Rh50-GAL4; UAS-mCD8::GFP* | Figures 1A, 1B, and 1D<br>Figure 3B<br>Figures S1B–S1D |
| *w; Amt-GAL4; UAS-mCD8::GFP* | Figure 1C<br>Figure 3C |
| *w; UAS-mCD8::GFP; Ir92a-GAL4* | Figure 1E |
| *w; Ir31a-GAL4; UAS-mCD8::GFP* | Figure 1F |
| *w; UAS-mCD8::GFP; Ir75d-GAL4* | Figure 1G |
| *w; 10X UAS-myc-APEX2-Orco; Or56a-GAL4*<br>*w; 10X UAS-myc-APEX2-Orco; Or47b-GAL4*<br>*w; 10X UAS-myc-APEX2-Orco, Or88a-GAL4; +*<br>*w; 10X UAS-myc-APEX2-Orco; Ir75c-GAL4*<br>*w; 10X UAS-myc-APEX2-Orco; Or47a-GAL4* | Figure 1I<br>Figures S1I and S1J |
| *w; Rh50-Gal4/repo-GAL80; Rh50-Gal4/UAS-mCD8::GFP* | Figures 1J and 1K |
| *w; UAS-GCaMP7s/Bl<sup>l</sup>; Rh50-GAL4/+* | Figure 2A |
| *w; UAS-GCaMP7s/Sp; Ir92a-GAL4/+* | Figure 2B |
| *w; UAS-DTA/Cyo; Ir92a-GAL4-1* | Figures 2C and 2D |
| *w; UAS-DTA/Cyo; +*<br>*w; UAS-DTA/Cyo; Rh50-GAL4* | Figures 2C and 2D<br>Figures 3L and 3M |
| *w; +; Rh50-GAL4* | Figures 3L and 3M |
| *w; Rh50-GAL4/+; UAS-GCaMP7s/+* | Figures 3A, 3E–3J |
| *w; Ir64a-GAL4; UAS-mCD8::GFP* | Figure 3D |
| *w; Ir25a-GAL4; UAS-mCD8::GFP* | Figure 4A |
| *w; UAS-mCD8::GFP; Ir76b-GAL4* | Figure 4B |
| *w; UAS-mCD8::GFP; Ir8a-GAL4* | Figure 4C |
| *w; UAS-mCD8::GFP; Orco-GAL4* | Figure 4D |
| *w; Ir25a<sup>2</sup>; +* | Figure 4E |
| *+; +; + (Canton-S, CS)* | Figure 4E<br>Figure S2<br>Figures S3B and S3E |
| *w; +; Rh50<sup>l</sup>* | Figure 4F<br>Figures S4B and S4C |
| *w; +; + (wCS)* | Figure 4F<br>Figures S3C and S3D<br>Figures S4B and S4C |
| *w<sup>1118</sup>; +; Amt<sup>l</sup>*<br>*w<sup>1118</sup>; +; + (isogenic background)* | Figure 4G |
| *w; UAS-GCamp6s; Ir75d-GAL4*<br>*w; UAS-GCamp6s/UAS-Amt; Ir75d-GAL4/+* | Figure 4H |
| *w; UAS-Amt; +*<br>*w; +; Ir75a-GAL4*<br>*w; UAS-Amt; Ir75a-GAL4* | Figure 4I |
| *w; +; Or22a-GAL4*<br>*w; UAS-Amt; Or22a-GAL4*<br>*w; UAS-Amt; +* | Figure 4J |
| *w; UAS-AgAmt; Ir75a-GAL4*<br>*w; UAS-AgAmt; +* | Figure 4K |
| *w; Ir41a-GAL4; UAS-mCD8::GFP* | Figure S1E |
| *w; UAS-mCD8::GFP; Or35a-GAL4* | Figure S1F |
| *w; Ir76a-GAL4; UAS-mCD8::GFP* | Figure S1G |
| *w; +; Ir92a<sup>l</sup>* | Figures S3C and S3D |
| *+; +; Ir92a<sup>l</sup>* | Figures S3B and S3E |

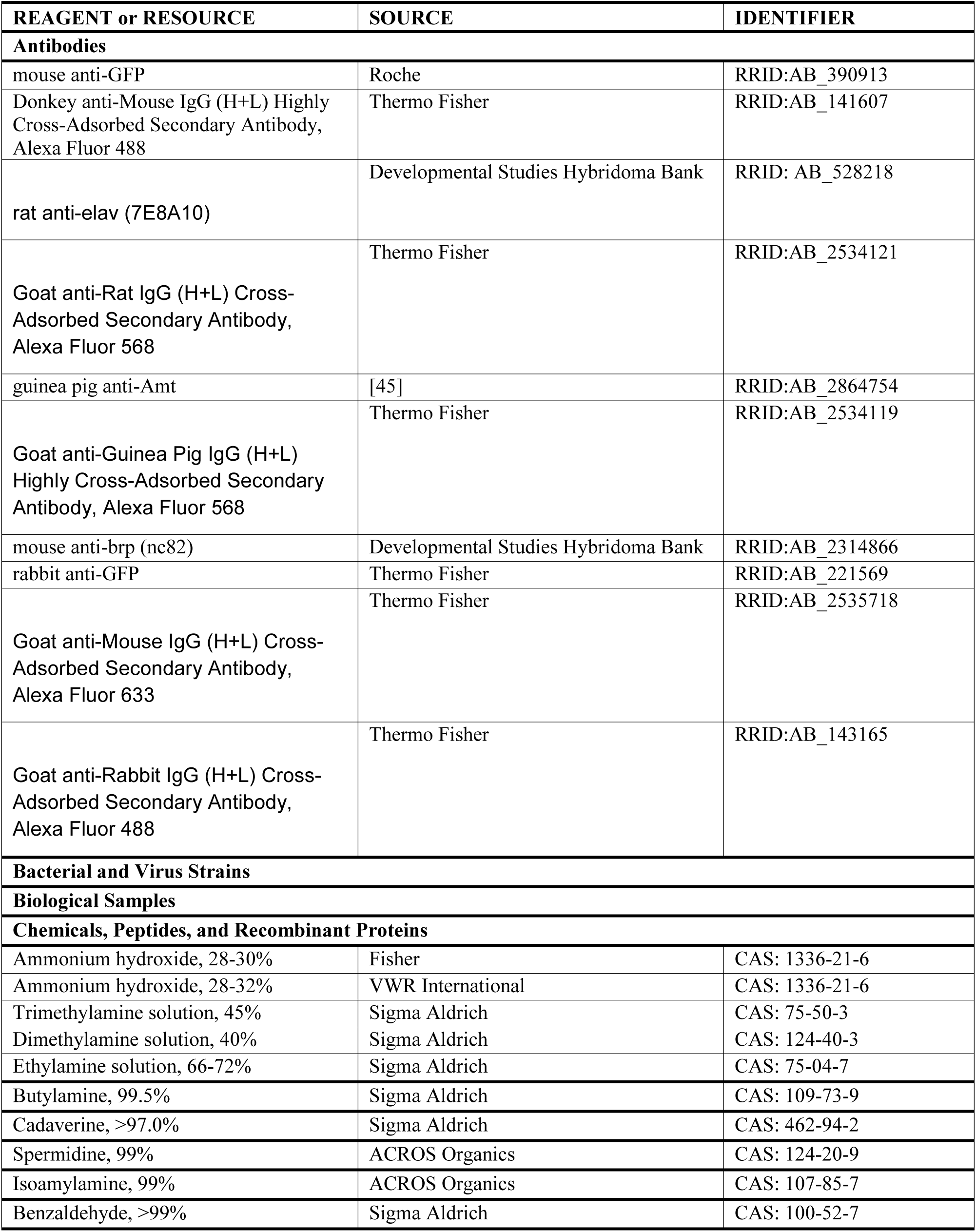
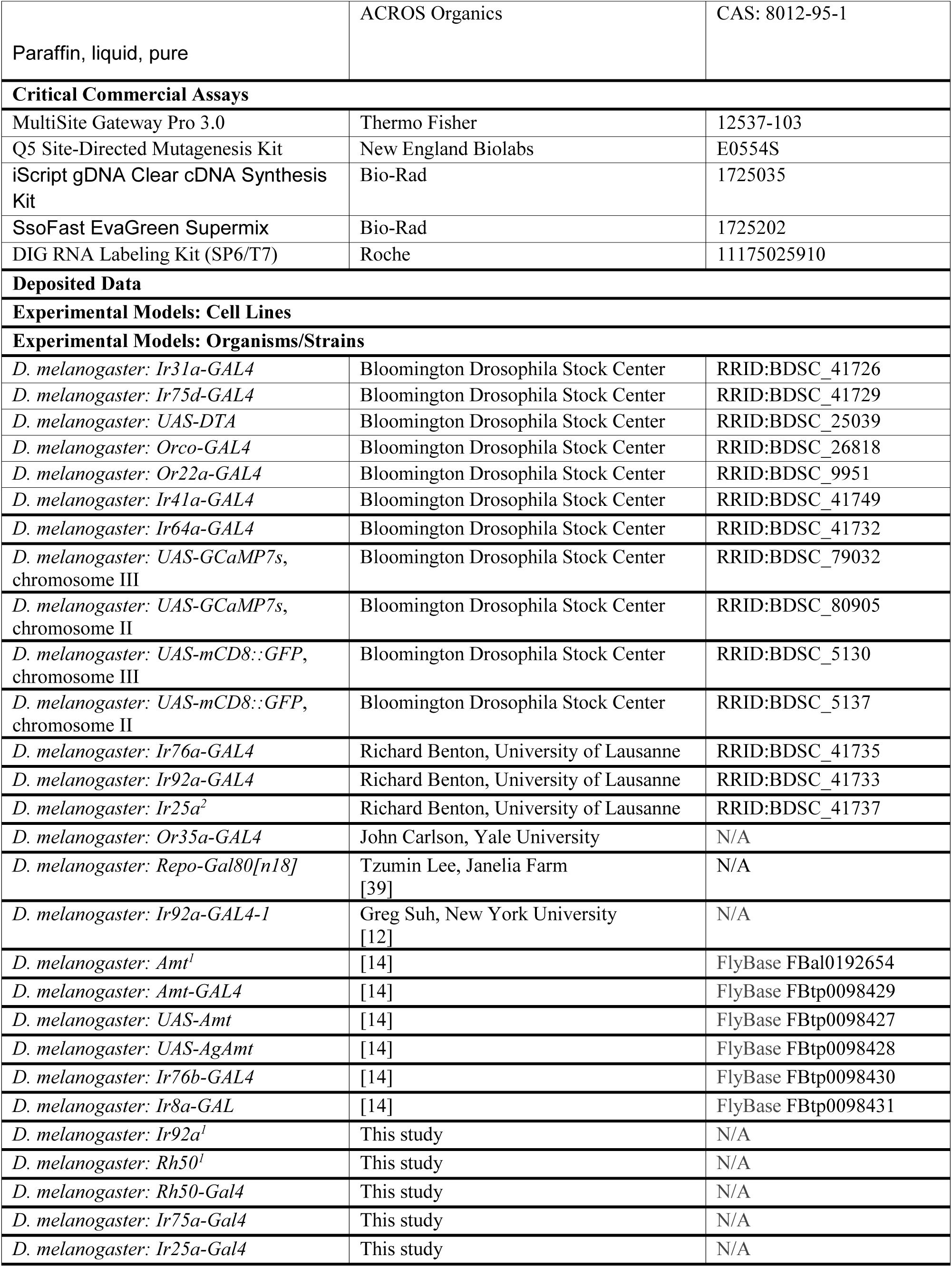
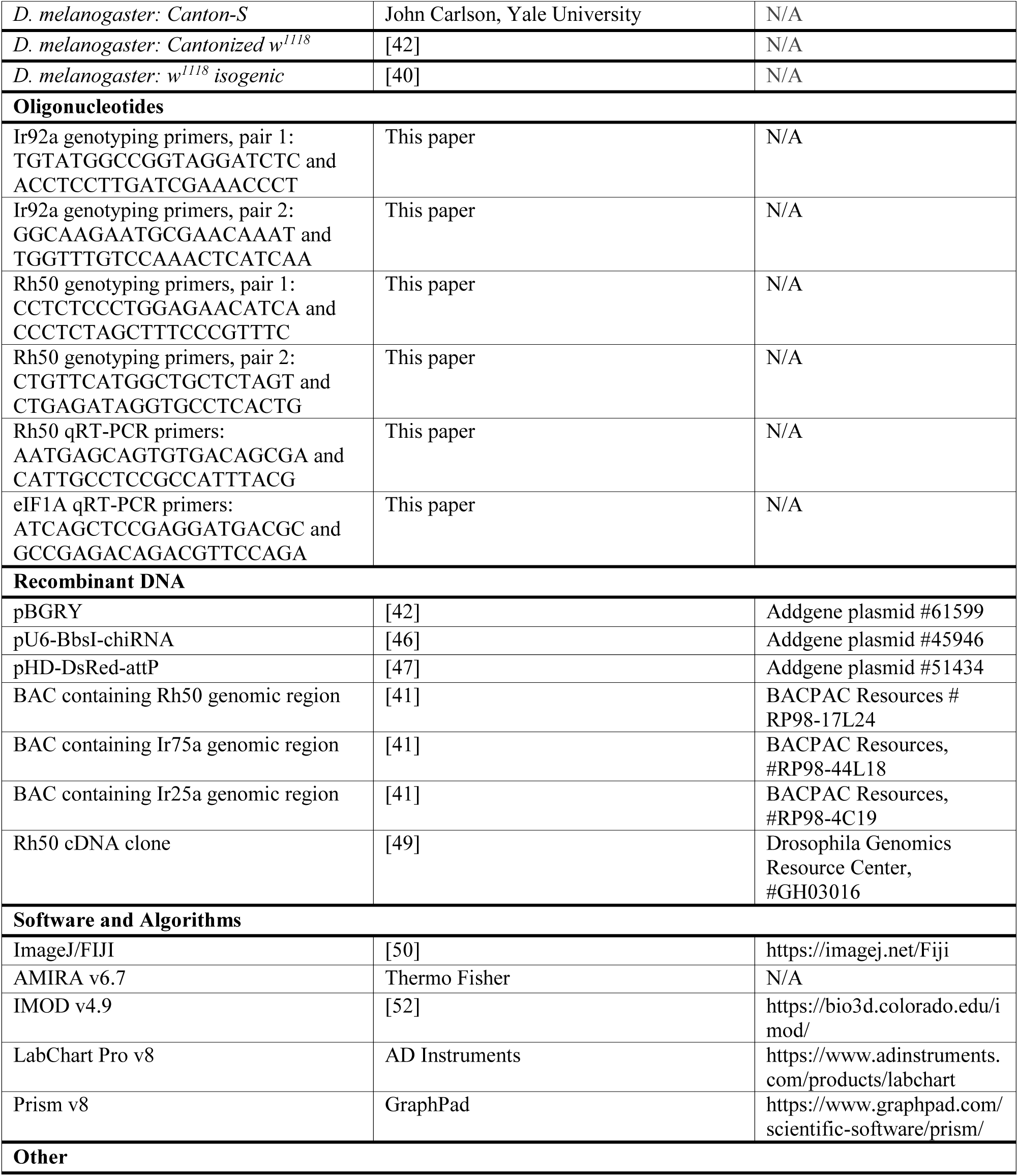
KEY RESOURCES TABLE.

